# Reinforcement learning derived chemotherapeutic schedules for robust patient-specific therapy

**DOI:** 10.1101/2021.04.23.441182

**Authors:** Brydon Eastman, Michelle Przedborski, Mohammad Kohandel

## Abstract

The in-silico development of a chemotherapeutic dosing schedule for treating cancer relies upon a parameterization of a particular tumour growth model to describe the dynamics of the cancer in response to the dose of the drug. In practice, it is often prohibitively difficult to ensure the validity of patient-specific parameterizations of these models for any particular patient. As a result, sensitivities to these particular parameters can result in therapeutic dosing schedules that are optimal in principle not performing well on particular patients. In this study, we demonstrate that chemotherapeutic dosing strategies learned via reinforcement learning methods are more robust to perturbations in patient-specific parameter values than those learned via classical optimal control methods. By training a reinforcement learning agent on mean-value parameters and allowing the agent periodic access to a more easily measurable metric, relative bone marrow density, for the purpose of optimizing dose schedule while reducing drug toxicity, we are able to develop drug dosing schedules that outperform schedules learned via classical optimal control methods, even when such methods are allowed to leverage the same bone marrow measurements.

## 1 Introduction

In mathematical models of cancer treatment, one is often concerned with finding a chemotherapeutic dosing schedule that is optimal in some capacity [1–3]. Each different model allows distinct forms of this optimality metric. This shaping of optimality metric will often involve a trade-off of some variety: incredibly high doses of a potent chemotherapeutic can certainly annihilate the cancerous cells in tissue but in so doing will largely cause a great deal of harm to the patient. Modelers, then, are concerned with mathematically formulating this optimality metric in a way that preserves the health and longevity of their patient (virtual or otherwise). For instance, in Ref. [2] the authors were concerned with maximizing the reduction in the total number of cancerous cells with the minimal total chemotherapeutic dose. They achieved this control by sampling 50 virtual patients from a particular parameter distribution, training 50 different reinforcement learning agents on differential equations representing the tumour growth of these patients, and applying these agents to these patients. In contrast, in Ref. [1] the authors concerned themselves with maximizing the chemotherapeutic dose while minimizing the damage to healthy, proxy cells in the bone marrow. Practically, these are two (similar and related) methods for achieving the same ends, but the particulars of their formalization can lead to drastically different qualitative results.

In any situation, the models used to represent the delivery of the chemotherapeutic and the associated reduction in cancer cells can inform the choice of optimality metric. So too can the choice of model inform the method by which a modeller can find such an optimal dosing schedule of a chemotherapeutic. One such method, as employed in Ref. [1], is that of optimal control theory. When the model that governs the behaviour we are trying to assert some control over is codified by differential equations, optimal control theory can provide a methodology for finding the dose delivery function that maximizes whatever optimality metric the modeller chooses (if such a metric has a maximum) [4]. In some situations, this optimal control can be accomplished analytically as in Ref. [1], in others (such as the objective functional presented in Eq. (6)) numerical techniques such as those employed by the GEKKO package may be required [5].

A reinforcement learning approach can also be employed to maximize a given optimality metric (see, for instance, Ref. [2]). In a reinforcement learning context, when the state of the model at a given time *t* is known, one can construct a controller function via the learned optimal policy. In contrast to optimal control, reinforcement learning more easily lends itself to situations where the model behaviour is governed by systems more complicated than just those that can be represented with differential equations (for instance, reinforcement learning has had great success in solving Atari games, arcade games, Backgammon, etc. [6–8]). In particular, as illustrated in Figure 1, a reinforcement learner need only be provided with the action space of the environment; all other details about the environment are effectively a black-box. The agent takes an action, the environment then changes as a result according to some rule-set the learner need not have access to, and a reward is issued. The reinforcement learner then evolves to maximize total cumulative reward, not just the immediate reward benefit. Importantly, the environment may be governed by a deterministic set of differential equations, by a stochastic agent-based model, or by rules entirely determined by data. In this capacity, the black-box nature of the reinforcement learner environment is enticing to applied mathematicians as it allows the capacity to perform numerical learning experiments in a regime that was previously untractable. Indeed, recent advancements in computing power have allowed for the tractability of model-free reinforcement learning [9]. With the advent of big data sets and quantitative medicine, reinforcement learning can be used to leverage real world data as well as deterministic, validated models in order to learn a control in complicated contexts. Presently, we consider the environment to be governed by a system of simple differential equations to establish a framework methodology that can, in the future, be extended to other, more realistic domains.

**Figure 1:**
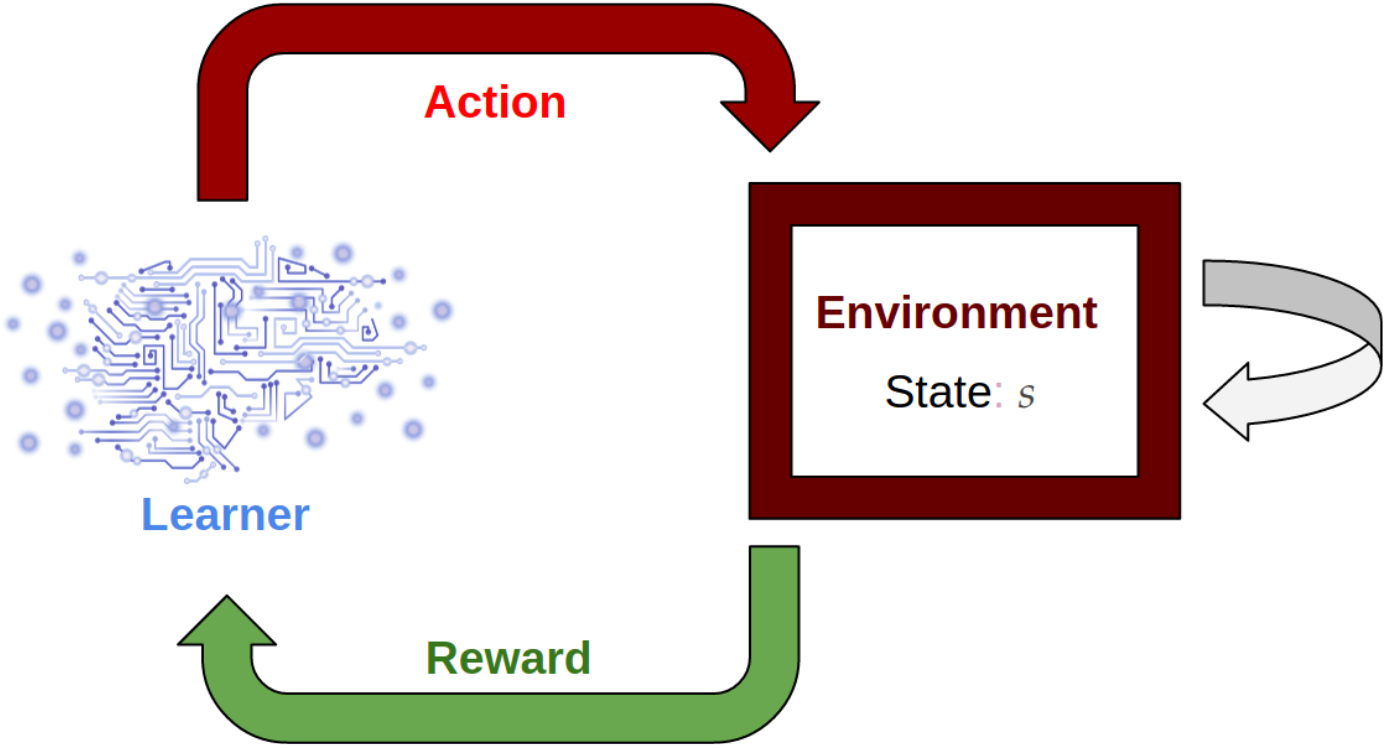
A reinforcement learning agent interacts with an environment as if the environment were a black-box. This process potentially changes the state of the environment and results in some reward for the learner. All that the learner needs to be provided with is the action space and a suitable reward function to determine an optimality metric.

In this study, as an example, we consider a simple differential equation model from Refs. [1,10–12]. This phenomenological model describes the growth of breast and ovarian solid tumors at a cellular level within a particular patient. The parameters of the model describe rates of cell-to-cell interaction and are incredibly difficult to measure in practice. In particular, the methods used in Refs. [1, 11] to parameterize this model only allow discovery of nominal, mean values of such parameters from multiple mouse models. While these parameters can help to capture the qualitative behaviour of the response of a tumour to a particular chemotherapeutic, the model can certainly not be considered to be a validated model in human cancers. However, even for a more robust, validated phenomenological model the issue of patient-specific parameter identification still remains. Whenever the parameter values used for these models are determined by population level data the modeller may not know, *a-priori*, the particular patient-specific parameters. In contexts where there is a demonstrable sensitivity to small perturbations in the parameter values there is a concern that the nominal parameters, and any chemotherapeutic control thereby derived, may not robustly describe the most optimal response for a particular parameterization. To that end, in this paper we explore how chemotherapeutic controls derived from mean value parameters can be used on models of patients with perturbed, unknown parameter values. In particular we leverage the power of deep double Q learning [6] to derive the chemotherapeutic control in a manner that provides learned dosing schedules that are robust to perturbations in parameter values in this sensitive system. Importantly, the reinforcement learning agent is unaware during training of the patient-specific parameter values on which it is evaluated, in contrast to Ref. [2] where multiple agents were trained on systems encoded by these parameter values exactly. In Ref. [1], the authors analytically derive the continuous optimal control of the tumor growth model used in this work under a particular objective functional. Here, we consider a similar optimal control problem but wherein both the drug dose and time are discretized.

The manuscript is organized as follows. In Section 2 we introduce the differential equation model and provide the mean parameter values that comprise the nominal virtual patient. We also provide an analytic characterization of the initial conditions used in the model simulations as a function of these parameter values in Section 2.1. In Section 2.2 we demonstrate that the model we consider is quite sensitive to local perturbations of the parameter values. In Section 3.1 we define the optimal control problem considered and the objective functional by which the optimal scores are deduced. In Section 3.2 we discuss the method by which virtual patients were created for testing and training purposes. In Section 3.3 we lay out the training process used for solving the reinforcement learning problem and the discrete optimal control problem and in Section 3.4 we discuss the hyperparameter tuning process of the deep double Q learning algorithm.

In Section 4.1 we compare the optimality of schedules learned by applying the reinforcement learning derived policy on unknown testing patients with that of the optimal schedule of the nominal patient being applied to these testing patients. In the former case, the relative bone marrow of these unknown patients was leveraged for the purpose of customizing the dosing schedule and reducing drug toxicity whereas, in the latter case, the same, nominal schedule is applied to all testing patients. In Section 4.2, we extend the optimal schedules to leverage relative bone marrow measurements as well by employing a version of nearest neighbour interpolation on the optimal control of various training patients, whose particular patient specific parameter values are treated as known, to personalize dose schedules for testing patients. This nearest testing neighbour optimal control is then compared with the previous reinforcement learning agent. Finally, we present a summary and discussion of our results in Section 5.

## 2 Tumor Growth Inhibition Model

We consider the two-compartment mathematical model of cell-cycle specific chemotherapy first introduced in Ref. [11], which is an extension of earlier work [10]. The model consists of a population of proliferating cells and a population of quiescent cells, where the time evolution of cell populations is depicted in Figure 2 and is governed by the following set of coupled ordinary differential equations:

**Figure 2:**
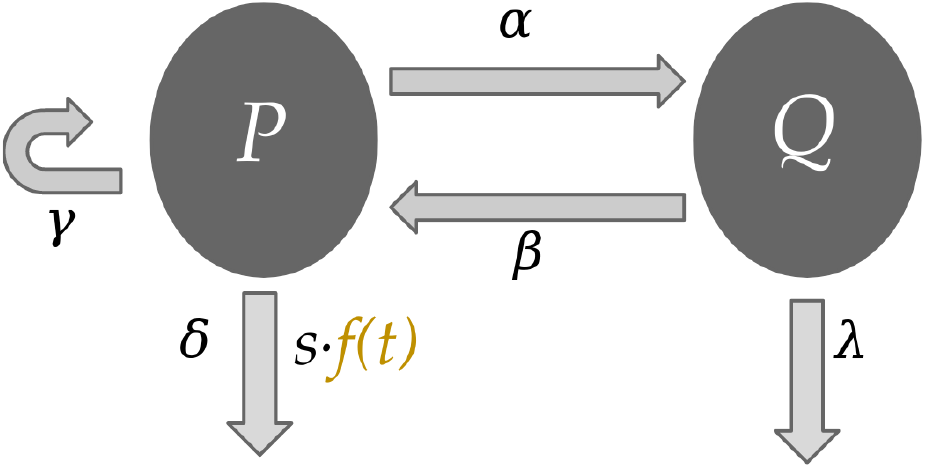
The two-compartment tumor growth inhibition model described by Eq. (1). Proliferative (*P*) cells and quiescent (*Q*) cells can both die naturally at the constant rates *δ* and *λ*, respectively. However, only proliferative cells can self-renew (at the constant rate *γ*) and be killed by the dose of a chemotherapeutic *f* (*t*). Moreover, proliferative cells are allowed to become quiescent (at constant rate *α*) and quiescent cells are allowed to become proliferative (at constant rate *β*).

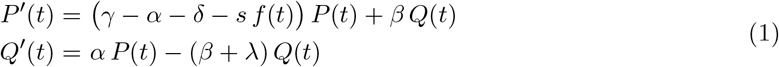

In the model, *P* (*t*) represents proliferative cells and *Q*(*t*) represents quiescent cells. The model captures the growth of proliferative cells at a constant rate *γ*, the transformation of proliferative cells into quiescent cells at a constant rate *α*, and the apoptosis of proliferative cells at a constant rate *δ*. Similarly, quiescent cells leave quiescence and become proliferative at a constant rate *β* and undergo apoptosis at a constant rate *λ*. The time-dependent function *f* (*t*) represents the dosing schedule of a chemotherapeutic where *s* represents the relative strength of the administration of such a chemotherapeutic. In particular, it is assumed that *f* (*t*) ∈ [0, 1]. While parameters *γ, δ, α, β*, and *λ* are patient-specific parameters depending on the nature of the disease being modelled, parameter *s* is a phenomenological hyper-parameter of the model describing the relative strength of the chemotherapeutic.

The proliferating cell compartment contains cells at each of the four phases of cell cycle (gap period G1, synthetic period S, second gap period G2, and mitosis M) to reduce the complexity of the cellular states. While resting cells are affected to a small extent by cell-cycle specific chemotherapy, the model (1) assumes that the chemotherapy *f* (*t*) affects only the proliferating cells. The model does not include details from other aspects of the patient’s context, notably it ignores the effects of age, sex, spatial information of the tumor, and any applicable comorbidities.

In Ref. [10] the authors parameterize Eq. (1) with values that are suitable for describing breast cancer and ovarian cancer, as determined by mouse models. In Ref. [1] the authors provide an additional parameter set for determining the effect of chemotherapy on healthy bone marrow cells. This allows one to, for a given chemotherapy dosing schedule *f* (*t*), model the effect of chemotherapy on both the healthy bone marrow cells and the malignant cancerous cells. Hence, by evolving two decoupled copies of Eq. (1), one parameterized with values corresponding to a particular cancer and the other with bone marrow parameter values, we can monitor the cancer-killing effects of a chemotherapeutic schedule and the associated chemotherapeutic toxicity in the patient. The parameter values are summarized in Table 1.

**Table 1:** Parameter values for breast cancer cells, ovarian cancer cells, and bone marrow cells as obtained from Refs. [1, 10], and values of 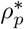 as determined by Eq. (3).

| **Bone marrow** |  |  |
| --- | --- | --- |
| Parameter | Nominal value | Units |
| $\gamma$ | 1.470 | days<sup>-1</sup> |
| $\delta$ | 0.000 | days<sup>-1</sup> |
| $\alpha$ | 5.643 | days<sup>-1</sup> |
| $\beta$ | 0.480 | days<sup>-1</sup> |
| $\lambda$ | 0.164 | days<sup>-1</sup> |
| $\rho_p^*$ | 0.103 | - |
| **Breast cancer** |  |  |
| Parameter | Nominal value | Units |
| $\gamma$ | 0.500 | days<sup>-1</sup> |
| $\delta$ | 0.477 | days<sup>-1</sup> |
| $\alpha$ | 0.218 | days<sup>-1</sup> |
| $\beta$ | 0.050 | days<sup>-1</sup> |
| $\lambda$ | 0.000 | days<sup>-1</sup> |
| $\rho_p^*$ | 0.200 | - |
| **Ovarian cancer** |  |  |
| Parameter | Nominal value | Units |
| $\gamma$ | 0.6685 | days<sup>-1</sup> |
| $\delta$ | 0.4597 | days<sup>-1</sup> |
| $\alpha$ | 0.2225 | days<sup>-1</sup> |
| $\beta$ | 0.0500 | days<sup>-1</sup> |
| $\lambda$ | 0.0000 | days<sup>-1</sup> |
| $\rho_p^*$ | 0.3600 | - |

### 2.1 Derivation of the Proliferative Fraction

We begin modelling the proliferative and quiescent components of Eq. 1 under the assumption that the tumour has evolved in the absence of any chemotherapeutic agent until a steady state, in terms of the proportion of these cells, has been reached. To that end, we define the proliferative ratio of the tumor at time *t* and the steady-state proliferative ratio as

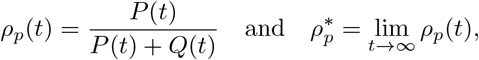

respectively. To analytically calculate the closed form solution of the steady-state proliferative ratio in the absence of a chemotherapeutic, we set *s* = 0 and consider

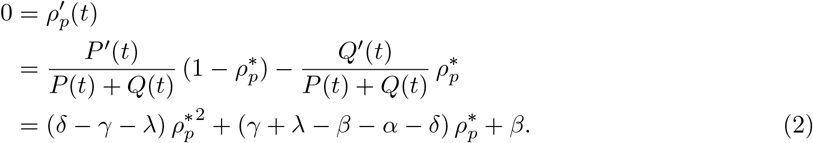

Thus, 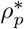 is a root of the quadratic in Eq. (2). For the parameter values presented in Table 1 the quadratic in Eq. (2) has only one positive (real) root, namely

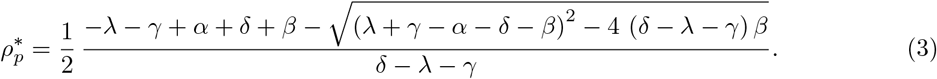

The values of 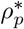 corresponding to the parameters for ovarian cancer, bone marrow, and breast cancer are included in Table 1. Thus the initial data for Eq. (1) considered in this study are given by 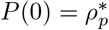 and 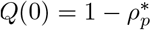.

### 2.2 Local Sensitivity Analysis

Here we investigate the sensitivity of the outputs for the tumor growth inhibition model, Eq. (1), to perturbations in the nominal parameter values of the model-specific parameters presented in Table 1. To compute the sensitivities, we change the values of the parameters *γ, δ, α, β*, and *λ* one-at-a-time by a small amount, Δ*p*. We take Δ*p* to be +1% of the nominal parameter value *p*_0_. Then the relative sensitivity of each model population *x* = ⟨*P*(*T*), *Q*(*T*)⟩ for the parameter is calculated as follows:

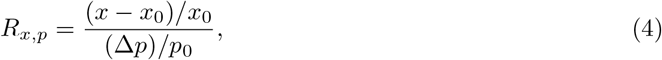

where subscripts denote nominal values. The initial conditions of the simulations were recalculated according to Eq. (3) for each perturbed parameter value and the simulations were run until *T* = 21 days. We plot the results in Figure 3.

**Figure 3:**
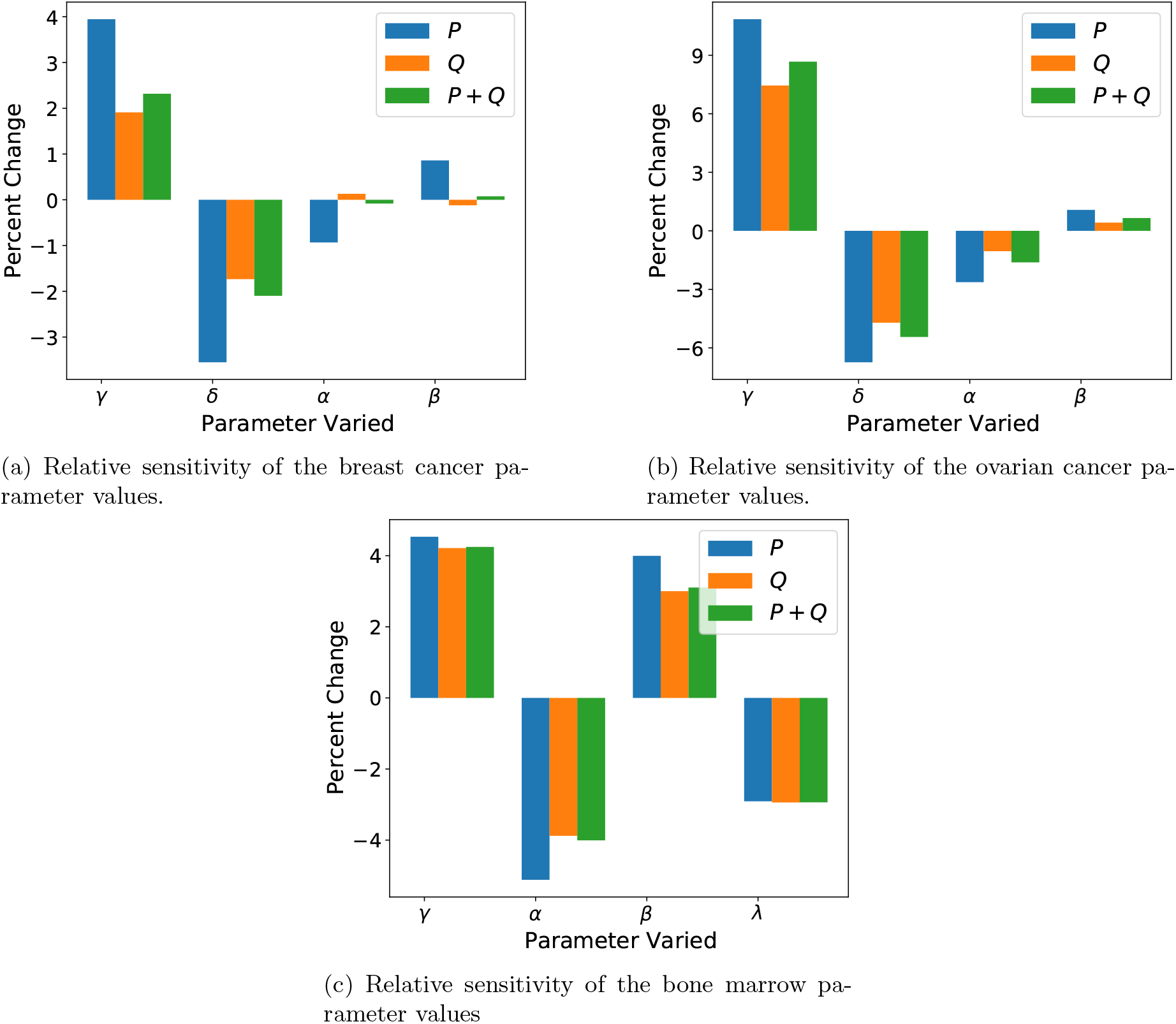
Relative sensitivity of Eq. (1) under the parameter sets from Table 1. Parameters with zero value (*δ* for bone marrow and *λ* for breast and ovarian cancer) were ignored and not displayed in this figure.

Importantly, the results of Figure 3 indicate that the model exhibits substantial sensitivity due to relatively small perturbations in the patient-specific parameter values. This type of sensitivity is common in models that experience regimes of exponential growth, which are common in cellular models of cancer [12]. For the parameter sets corresponding to breast cancer, Figure 3a indicates a mean (absolute) change of roughly 2.3% in *P* cells, 0.97% in *Q* cells, or 1.14% in all cell types given a 1% perturbation to a singular parameter. For ovarian cancer the model is even more sensitive, demonstrating a mean (absolute) change of roughly 5.3% in *P* cells, 3.4% in *Q* cells, or 4.1% in all cell types given a 1% perturbation to a singular parameter. For bone marrow, similar extreme sensitivities are observed with a mean (absolute) change of roughly 4.1% in *P* cells, 3.5% in *Q* cells, or 3.6% across all cell types. Importantly, Figure 3 demonstrates that even for the least sensitive parameter set (the breast cancer parameter set), small perturbations to individual parameters can still elicit large differences in the evolution of a tumour if one is unlucky enough that such a perturbation occurred in either *γ* or *δ* (the self-renewal and death rate of proliferative cells, respectively).

## 3 Methods

### 3.1 Chemotherapeutic Control

To determine the optimal chemotherapeutic control, we follow Ref. [1] and introduce an objective functional with the form

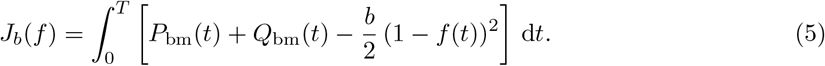

Maximizing this objective functional enables the derivation of an optimal chemotherapy dosing schedule of duration *T* days for a particular patient. In the notation of Eq. (5), *P*_bm_ and *Q*_bm_ refer to the proliferative and quiescent compartments of Eq. (1) parameterized to describe the behaviour of bone marrow. In effect, this leads to a chemotherapeutic schedule that biases the optimizer toward applying a larger dose of chemotherapeutic, as governed by *f* (*t*), while also maximizing the total number of bone marrow cells in the patient (to reduce drug toxicity). The non-negative hyperparameter *b* is then a scaling factor representing the relative importance of these two mechanisms. If *b* ≫ 1, then delivering the largest chemotherapeutic dose possible is the most desirable action for the optimizer, even at the cost of decimating the bone marrow cell count. Contrarily, if 0 ≤ *b* ≪ 1 then the optimizer is biased toward preserving bone marrow even at the cost of lower cancer kill. Plots of such a chemotherapeutic dosing function *f*, obtained via the analytical method for deriving the continuous optimal control as in Ref. [1], for various *b* values are presented in Figure 4.

**Figure 4:**
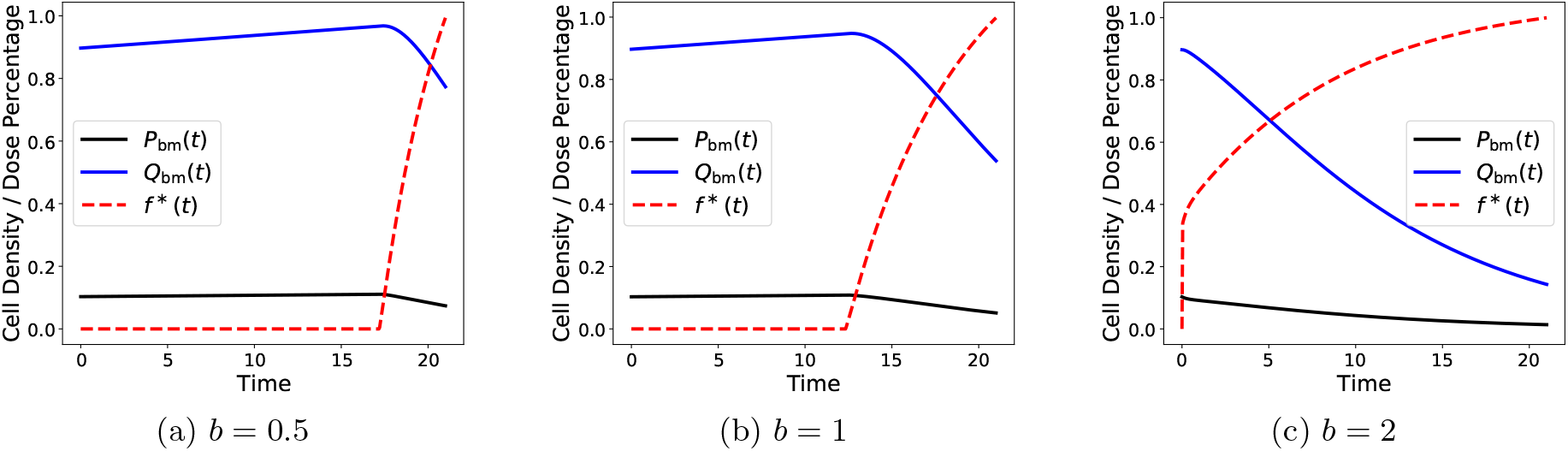
A plot of the proliferative cell proportion (black), the quiescent cell proportion (blue), and the optimal chemotherapeutic control *f* ^*\**^(*t*) (dashed red) for different values of *b*. The objective functional used to achieve this optimal control is given via Eq. (5). Small values of *b* correspond to weighting preservation of the bone marrow as more important and larger values of *b* correspond to weighting total drug delivery as more important.

In this work, the particular functional form of Eq. (5) is not of primary concern. Certainly other forms could be suggested to achieve similar qualitative goals. For instance, consider the functional

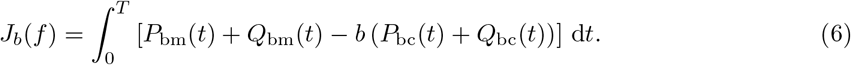

In the notation of Eq. (6), *P*_bc_ and *Q*_bc_ refer to the proliferative and quiescent compartments of Eq. (1) parameterized to describe the behaviour of solid breast cancer tumors. Hence, the functional in Eq. (6) describes the minimization of breast cancer cells while preserving the healthy, bone marrow cells. In this functional, the dependence on the chemotherapeutic dosing schedule *f* is implicitly included in the trajectories of *P*_bm_, *Q*_bm_, *P*_bc_, and *Q*_bc_. In any case, the exact formulation of this objective functional is an incredibly important choice for any modeller in a clinical context as it determines the metric by which the control is considered maximal and is outside the scope of this report.

While there are many methods in the field of optimal control theory that provide a methodology for obtaining such schedules, one could also employ techniques from reinforcement learning to discover chemotherapeutic dosing schedules. For instance, for a time *t* given the state vector

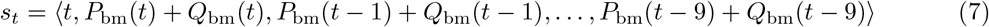

and chemotherapeutic dose *a*_*t*_ ∈ [0, 1], we define the immediate reward function as

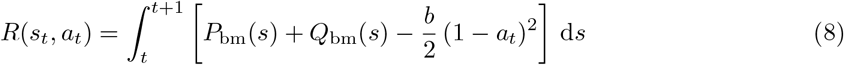

in order to elicit an analogous response in the reinforcement learner as achieved by the objective functional in Eq. (5). Explicitly, 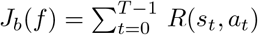, where *J*_*b*_ is given in Eq. (5) for appropriate piecewise constant functions *f*. To proceed, we use the reward function in Eq. (8) in the following form of the Bellman equations (see, for instance, Ref. [13])

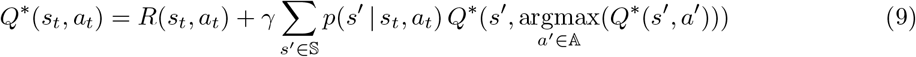

to derive an optimal policy as defined by

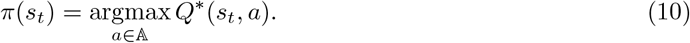

This policy can then be used to derive an optimal chemotherapy dosing schedule according to

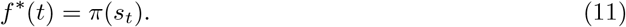

We provide a brief explanation of Equations 9-11 but direct readers to a more thorough source such as Ref. [13] for full details. As mentioned earlier, *R*(*s*_*t*_, *a*_*t*_) represents the immediate reward an agent receives for performing action *a*_*t*_ while in state *s*_*t*_. In contrast, *Q*^*\**^(*s*_*t*_, *a*_*t*_) represents a valuation of performing action *a*_*t*_ while in state *s*_*t*_. Notably, *Q*^*\**^(*s*_*t*_, *a*_*t*_) encodes the immediate reward *R*(*s*_*t*_, *a*_*t*_), but also encodes the discounted future rewards. Similarly, for a given *Q*^*\**^ function the policy *π*(*s*_*t*_) describes the optimal action to perform in a given state *s*_*t*_. As a result, *π*(*s*_*t*_) chooses an action *a*_*t*_ to maximize *Q*^*\**^(*s*_*t*_, *a*_*t*_) in a global manner as compared to the local process of choosing *a*_*t*_ to maximize immediate reward *R*(*s*_*t*_, *a*_*t*_). As such, the maximal action in a given state, as valued by *Q*^*\**^, may be one for which the payoff is not immediately obvious for multiple timesteps. The factor *γ* in Eqn. 9 is the discount factor of future rewards. The parameter *γ* is taken such that *γ* ∈ [0, 1] where *γ* = 0 corresponds to an agent that is focused on maximizing the immediate reward of their action and *γ* = 1 corresponds to an agent more concerned with increasing future reward than immediate. In general, a model describing a reinforcement learning environment may not be deterministic. In that regard, *p*(*s*′ |*s*_*t*_, *a*_*t*_) corresponds to the probability of ending up in state *s*′ after taking action *a*_*t*_ in state *s*_*t*_. For the model in Eqn. 1, no such stochasticity exists. As such, it is assumed that *p*(*s*′ |*s*_*t*_, *a*_*t*_) = 1 for exactly one *s*′ ∈ 𝕊 (namely *s*′ = *s*_*t*+1_). One can derive the drug dosing schedule *f* ^*\**^(*t*) as in Eqn. 11 by observing the state in some manner and then evaluating the policy at this state. For the case of the nominal patient, where the patient specific parameters are known, observing the state is as simple as integrating Eqn. 1. For an already trained model, one would construct the state vector (Eqn. 7) for a patient (virtual or otherwise) and then evaluate the policy at this state.

For a continuous time reinforcement learning agent, the optimal dosing schedule learned by this process for a given parameterization of Eq. (1) is identical to that derived via optimal control theory for that same parameterization, as in Ref. [1]. Of particular importance, however, is that as Eq. (11) demonstrates, once a policy has been learned, one can derive an optimal chemotherapy schedule by merely evaluating the policy at the state. Importantly, the state one evaluates the policy at need not be a state seen during the learning process. Indeed, in our study we concern ourselves with training the reinforcement learning agent on only the nominal virtual patient and developing chemotherapy schedules for 200 different testing virtual patients. By leveraging state vector information from these 200 different testing virtual patients (patients which encode an environment over which the agent has not trained) the reinforcement learning agent is able to personalize the dose delivery function. As a result, it is important that we define our state vector as something that is both practically measurable and phenomenologically linked to the objective functional we wish to optimize. As indicated by Eq. (7), when deriving the optimal chemotherapy dosing schedule according to Eq. (11), we are passing the optimal policy a 10-day window of measurements corresponding to bone marrow count relative to a time before treatment began as well as the current day of treatment (in order to satisfy the Markov property).

### 3.2 Perturbed Virtual Patients

We generate sets of virtual patients according to the following strategy. We consider parameter values from Table 1 and construct virtual patients by perturbing these parameter values. These perturbations are performed by scaling the mean parameter values in Table 1 by factors sampled from the space [1 − *k*, 1 + *k*] uniformly with Latin hypercube sampling [14]. Latin hypercube sampling, a space filling technique for drawing random samples, is especially important when the number of samples drawn is small in comparison to the size of the sample space and when the parameters of interest are uncorrelated. Given the phenomenological nature of the five parameters, *γ, δ, α, β*, and *λ*, we can assert that these parameters are uncorrelated. In particular, modifying any one of these parameters will create a distinct system under Eq. (1) that cannot be recovered by modifications to any number of the remaining parameters.

In this work, we generated virtual patients at perturbation levels of *k* = 0.15, 0.20, and 0.25, where *k* corresponds to the percent-change strength of perturbation. We generated six total sets of virtual patients: for training purposes, we generated 50 virtual patients at the 15%, 20%, and 25% perturbation strength level; for testing purposes, we generated 200 virtual patients at the 15%, 20%, and 25% perturbation strength level. The reinforcement learning agent was only trained on the nominal virtual patient and not virtual patients from the training or testing sets. The training virtual patients were utilized for the crafting of an optimal control strategy described in Section 4. In this regard, the testing virtual patients serve as a metaphor for the unknown patient-specific parameters of any particular patient in clinic. Both the testing and training virtual patients, for the non-zero parameters *γ, α, β*, and *λ*, are visualized in Figure 11.

### 3.3 Training Process

We numerically solve the Bellman equation, Eq. (9), by employing neural networks as in the deep double Q-learning algorithm [6]. In particular, we represent the *Q* function from the Bellman equation, Eq. (9), as a neural network. As a result, after training the network, the specific form of *Q* is the same for each testing virtual patient. However, as in Eq. (11), by supplying the bone marrow measurements for the testing virtual patient, the network can produce a personalized dose schedule that is different for each virtual patient. In terms of the architecture of the *Q* network, we take the network to have 10 inputs neurons (as determined by the length of the state vector), hd_1_ neurons in the first hidden layer, hd_2_ neurons in the second hidden layer, and 11 neurons in the output layers (corresponding to dose strength range from 0 to 1 inclusive in 0.1 increments). Each neural layer is activated with a rectified linear unit. We use batch-learning with a batch size of `bs` to minimize the mean squared error between the right and left hand sides of Eqn. 9 via an Adam optimizer with learning rate *α*. In Section 3.4 we discuss how we decide upon the values of these hyperparameters and list the particular values in Table 2. To train the network for a given hyperparameter set, we first run 5,000 exploratory time steps of the simulation performing random actions in order to fill the experience replay buffer. After every 21 time steps, the environment is reset by returning the differential equation model, Eq. (1), back to its initial conditions (as dictated by Eq. (3)). In particular, the initial state for the reinforcement learning algorithm is a length eleven vector where the first entry is a 0 and the remainder are unit entries (as the Eq. (3) initial conditions sum to 1). This window length of 10 for the bone marrow measurements was chosen empirically, as in Ref. [2]. For each time step of the simulation we choose a chemotherapy dose according to our network via an *ϵ*-greedy algorithm. We anneal *ϵ* linearly from *ϵ* = 1 to *ϵ* = 0.01 over 25,000 time steps. The *E*-greedy algorithm was only implemented during training, i.e. during evaluation the policy selection is deterministic as in Eqn. (10). After selecting a dose *a* ∈ 𝔸 = {0, 0.1, …, 1}, we apply the chemotherapy dose to the patient by holding *f* (*t*) = *a* constant over the timestep and evolving Eq. (1) (as such, we discretize not only in dose but in time as well). Next, we record a tuple of the state, action, reward, new state values. We then select a random batch of previously observed tuples and use them to approximate the right hand side of the Bellman equation, Eq. (9), in order to obtain a target for training the network.

**Table 2:** Hyperparameter values for the learning process from Section 3.3 as determined by the Bayesian optimizer from Section 3.4.

| Hyperparameter Values |  |  |
| --- | --- | --- |
| Parameter | Value | Description |
| $\mathbf{hd}_1$ | 64 | Dimension of first hidden layer in the neural network |
| $\mathbf{hd}_2$ | 96 | Dimension of second hidden layer in the neural network |
| $\gamma$ | 0.9553 | Discount factor from the Bellman equation (9) |
| $\alpha$ | 0.003809 | Learning rate for the Adam optimizer |
| $\mathbf{bs}$ | 96 | Batch size for the Adam optimizer |

This process is eventually stopped if the differential equation environment has been reset 50,000 times or if the best reward has not improved over the last 500 epochs. This constitutes one training run of the system. We perform 5 such training runs recording the run that achieved the objective functional score under Eqn. 5.

### 3.4 Hyperparameter Tuning

While our system has many model specific parameters, there are also a number of hyperparameters introduced during the training process. These are the learning rate for the Adam optimizer (*α*), the dimension of the two hidden layers (hd_1_ and `hd`_2_, respectively), the discount rate *γ* in the Bellman equation (Eq. (9)), and the batch size of the Adam optimizer used for learning (`bs`). In order to ensure optimal convergence and stability of the resultant networks, we must carefully select these values. For a single set of these five hyper-parameters we must execute the entire training process over again. Such a process is computationally extensive rendering a brute-force grid-search approach to hyperparameter optimization unfeasible. To that end, we instead use Bayesian optimization to explore this five-dimensional hyperparameter space more efficiently. We allow our Bayesian optimizer to sample 100 such hyperparameter samples from this hyperparameter space and perform training process described in Section 3.3 for each hyperparameter set. The Bayesian optimizer chose hd_1_ and `hd`_2_ from the set {64, 96, …, 256}, `bs` from the set {32, 64, …, 128}, *α* from the interval (10^−4^, 10^−1^), and *γ* from the interval (0, 1).

In Figure 5 we see the distribution of the objective functional score under Eqn. 5 for the 100 reinforcement learning agents under this hyperparameter tuning process. In particular, we note the cluster of 36 agents that converged to the network architecture with the theoretical maximal objective functional value, as determined by running a discretised version of the optimal control problem from Ref. [1] with the APOPT algorithm (as implemented by GEKKO [5, 15]). For a point of comparison, we calculated the expected score achievable by a random agent by calculating the mean value of the score obtained in 1,000,000 simulations where a dose from 0, {0.1, …, 1.0} was uniformly selected at each time step. This resulted in a mean objective functional value of 0.6806 with a standard deviation of the mean of 0.04811.

**Figure 5:**
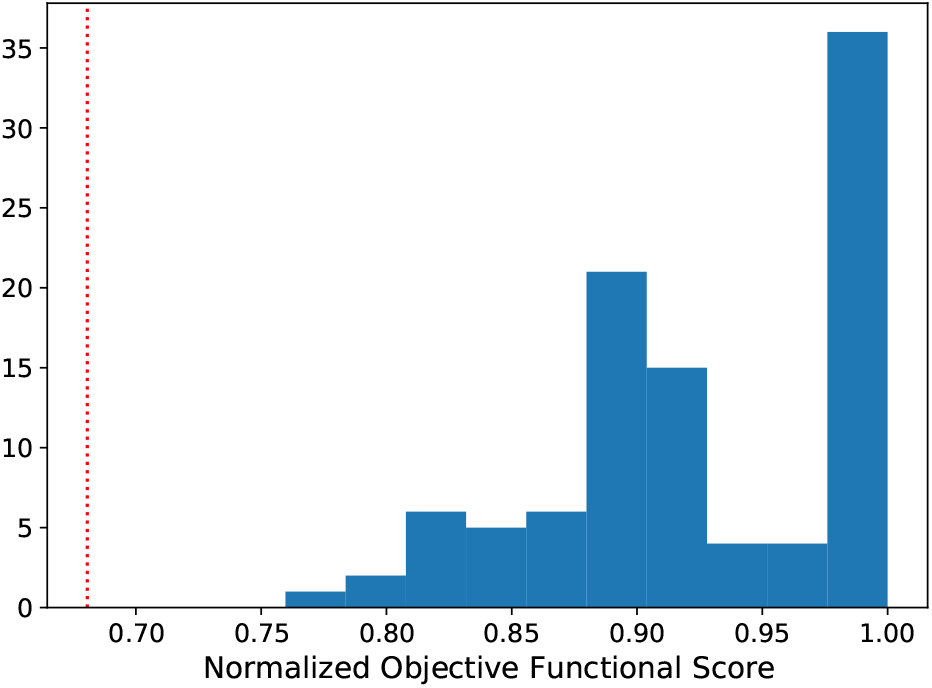
A histogram demonstrating all the scores obtained via the reinforcement learning process. The red dotted line indicates the expected score of a random agent.

To ascertain the identifiability and stability of these parameters, we consider the distribution of parameters that result in such objective functional value. To that end, in Figure 6 we consider the distribution of each individual hyperparameter and contrast this with the distribution of such hyperparameters from the agents that converged to network architecture that achieve an objective functional value value within 5% of the maximal possible reward. Importantly, we recognize that of the five hyperparameters, there is not a tight distribution after conditioning on objective functional value score. In fact, only the discount factor *γ* produces a conditioned distribution that is statistically different than the un-conditioned distribution (two-sample Kolmogorov-Smirnov p-value of approximately 0.001 [16]). This suggests that the particular values of the size of the hidden dimensions, learning rate, and batch size are not terribly sensitive parameters for the training of this reinforcement learning agent.

**Figure 6:**
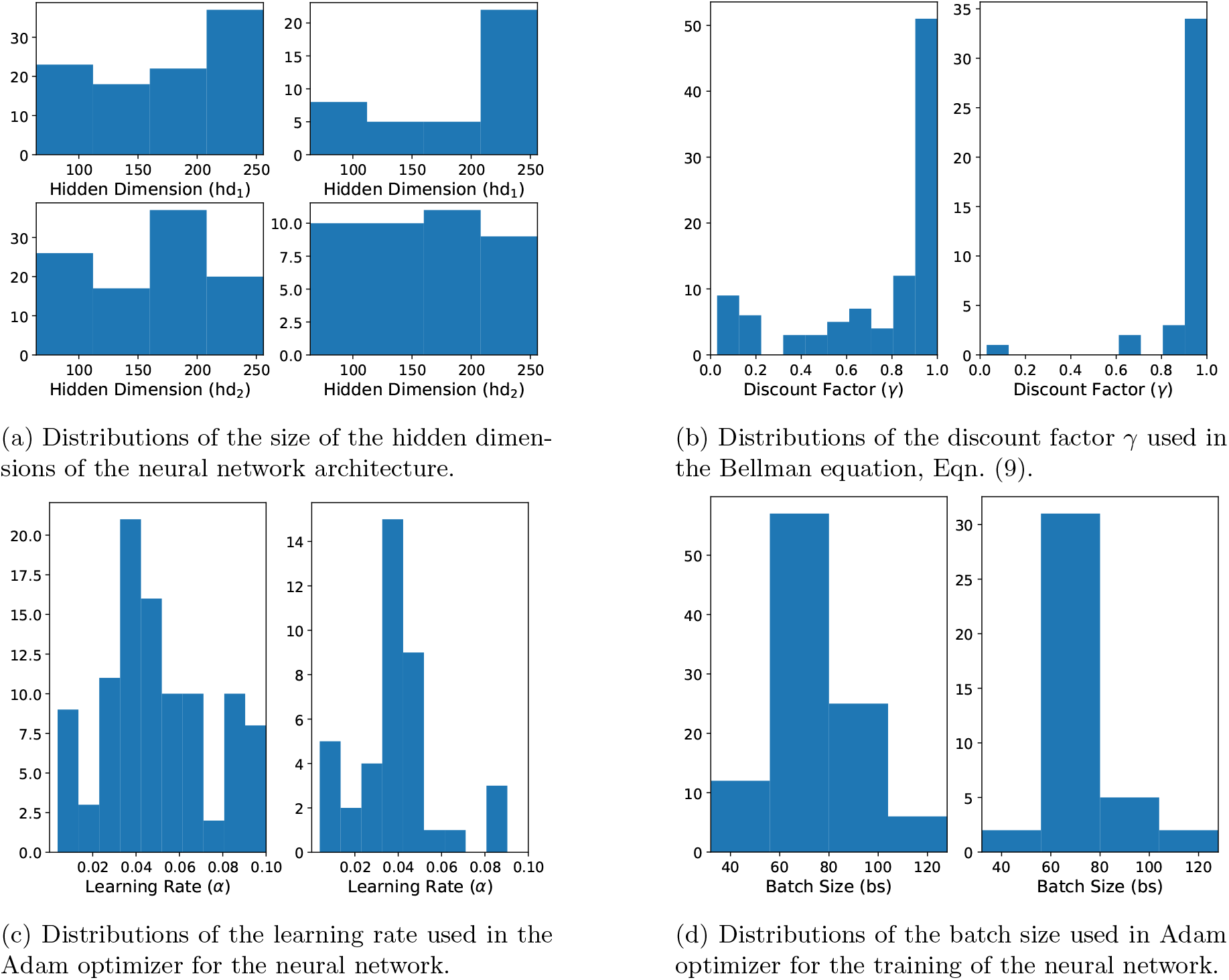
In all figures, the distributions on the left represent the total distribution of the hyperparameter explored by the Bayesian hyperparameter optimizer. In contrast, the distributions on the right in each figure indicate the distribution of the hyperparameter conditioned on the objective functional value being within 5% of the maximal theoretical score.

The Bayesian optimizer determined an optimal hyperparameter choice of hd_1_ = 64, `hd`_2_ = 96, *γ* = 0.9553, *α* = 0.003809, and `bs` = 96. Though, as the above discussion demonstrates, it is only the choice of *γ* that appeared to have any particularly strong impact on the convergence of the training process. A *γ* value close to 1 can be interpreted as representing an agent with a far horizon [13]. In particular, such an agent is less concerned with the immediate reward of a particular action and more concerned with the long-term, cumulative reward obtained by maximizing Eqn. 5 over all time.

## 4 Results

### 4.1 Contrasting a nominal reinforcement learning agent with a nominal optimal controller

We first begin by training a reinforcement learner on the nominal parameter set from Table 1 as per the method described in Section 3.3 using the hyperparameters for the training method from Table 2. During training, the reinforcement learning agent only interacted with the environment from Eq. (1) parameterized by the nominal set. Similarly, as a point of comparison, we used the APOPT algorithm from the GEKKO Python library to solve the discretized optimal control problem on the nominal parameter set [5, 15]. These two agents, one a reinforcement learning agent and the other a traditional optimal controller, were kept blind to the testing and training virtual patients. We then applied chemotherapeutic dosing schedules derived from both methods on the 600 testing virtual patients (200 virtual patients each at the 15%, 20%, and 25% perturbation strength level). In order to normalize the scores of these trials, we separately solved the discretized optimal control problem on these testing virtual patients using the APOPT algorithm from GEKKO again. Importantly, these 600 solutions were only used to ascertain the maximal possible objective functional value in order to scale the result of the blind agents. Finally, we compared the blind agents results by applying their chemotherapy derived strategies to the testing virtual patients, scaling the output according to the previously ascertained maximal possible reward. The results of this are presented in Figure 7 for the 3 different perturbation strength levels.

**Figure 7:**
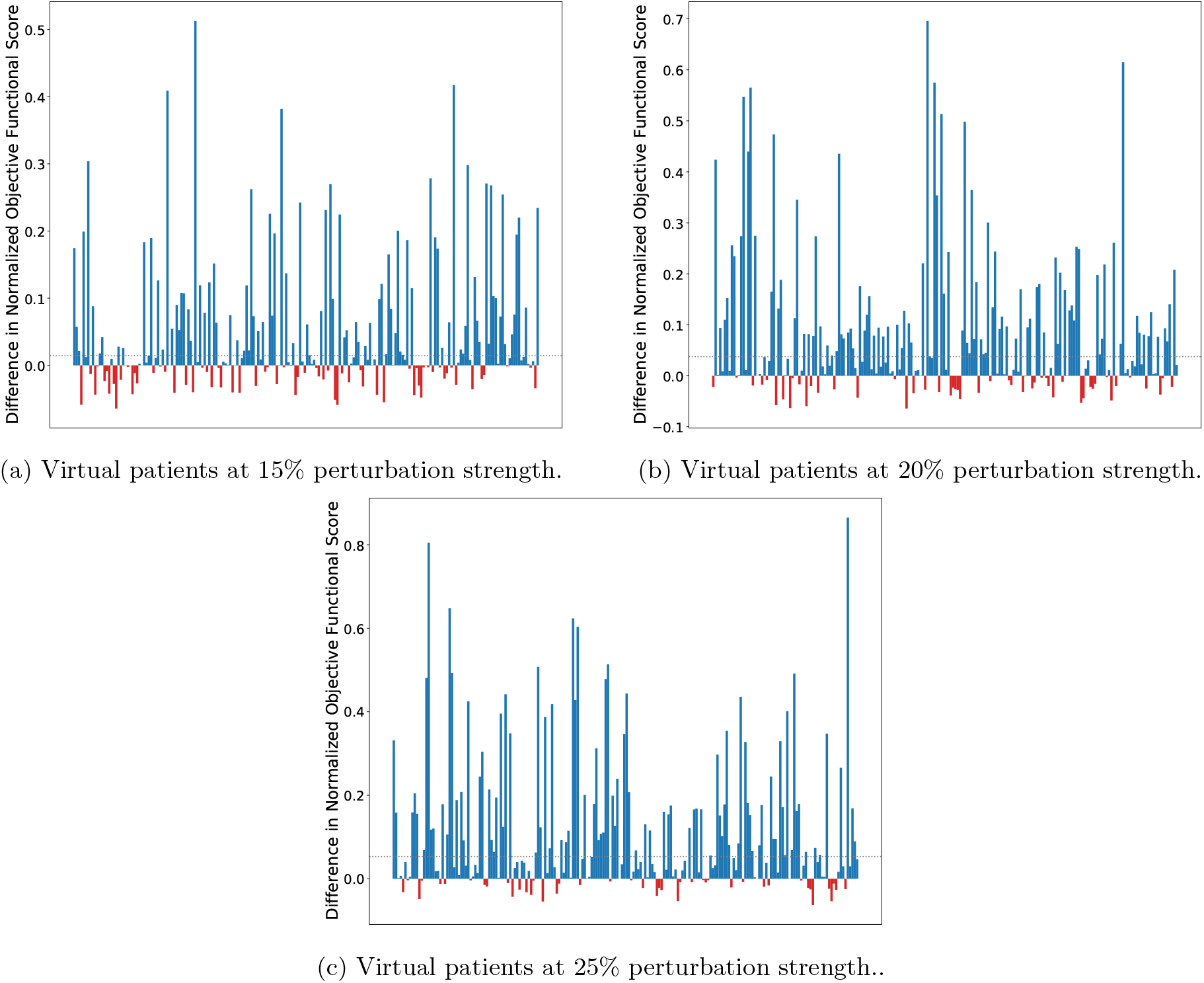
Bar plots of the difference between the scores obtained by the reinforcement learner derived policy and the scores obtained by the optimal control derived policy on all test virtual patients. Testing patients where the reinforcement learner outperformed the optimal controller are marked in blue and patients where the optimal controller outperformed the reinforcement learner are marked in red. The dotted grey lines in each plot indicate the difference of the median normalized scores of the reinforcement learner and the optimal controller.

Notably, the chemotherapy dosing schedule determined via optimal control for the nominal parameter set is a particular function *f* ^*\**^ that is the same for each virtual patient. In effect, each virtual patient is treated with the therapy schedule that is optimal for the mean-valued patient. In contrast, in the reinforcement learning derived schedule, the policy from Eq. (10) is the same for each virtual patient, but that policy is being fed a 10 day window of relative bone marrow cell counts from each virtual patient as a state vector. In effect, we are allowing the reinforcement learner to refine its dosing schedule given this information. By doing so, we are able to acquire a dosing schedule that is more robust to perturbations in these unknown, assumed to be unmeasurable, patient-specific model parameters by allowing refinements to be made based on a more easily measurable aggregate metric. Importantly, the state vector for the reinforcement learner is the sum of the *P*_bm_ and *Q*_bm_ compartments of Eq. (1) at discrete time points (in this case, daily) and not the individual measurements of *P*_bm_ and *Q*_bm_ separately. Given the scaling of Eq. (1) under the initial conditions from Eq. (3), these measurements are taken relative to the bone marrow mass prior to treatment, and so absolute measurements are not required. See Figure 13 for a visualization of these schedules on different virtual patients.

To quantify the performance differences of the two dose schedule processes over the testing virtual patients, we compared the distributions of scores with the non-parametric one-sided Wilcoxon signed-rank test [17]. For the cases represented in Figures 7a, 7b, and 7c we considered the alternative hypothesis to be that the median normalized score obtained by the reinforcement learner is larger than the median normalized score obtained by the optimal controller. We found at the 15% perturbation strength level a Wilcoxon statistic of 15103 corresponding to a *p*-value on the order of 10^−10^, at the 20% perturbation strength level we found a Wilcoxon statistic of 16875 corresponding to a *p*-value below machine precision, and at the 25% perturbation strength level we found a Wilcoxon statistic of 17815 corresponding to a *p*-value below machine precision. In all cases, we reject the null hypothesis and conclude that the reinforcement learning agent produces chemotherapeutic schedules with a higher median score on perturbed patients than the optimal controller. We notice that as the perturbation strength increases, the difference in the median and mean normalized scores increases as well from a difference in medians of 0.014 in the 15% case (difference of means of 0.052) to a difference in medians of 0.053 (difference of means of 0.116) in the 25% case. Hence, as the strength of the perturbation increases over this range, the reinforcement learner outperforms the optimal controller even further.

In Figure 8 we present the histograms of the normalized scores for each blind agent on the 600 different virtual patients. The histograms are semi-transparent in order to aid comparison where the blue colour represents the reinforcement learning agent and the orange colour the APOPT derived optimal controller agent. We note that the reinforcement learning agent has a large cluster of treatments in the 97.5% – 100% optimal bin (167 out of 200 in the 15% case, 158 out of 200 in the 20% case, and 167 out of 200 in the 25% case) and all treatments fall within 7.5% of the theoretical maximum. These scores are achieved without training on these virtual patients directly. In contrast, the optimal controller derived treatment is much more diffuse. As the strength of perturbation increases, the average score of the reinforcement agent derived schedule remains within 1.2% of optimum, while the average optimal control derived score decreases dramatically from 0.934 in the 15% case, to 0.900 in the 20% case, and finally to 0.873 in the 25% case. In particular, this suggests that the increase in performance of the reinforcement learner as a result of perturbation strength is due to the reinforcement learning agents’ capacity to remain non-sensitive to these perturbations, in contrast to the sensitivity seen in the schedules derived by the optimal controlling agent.

**Figure 8:**
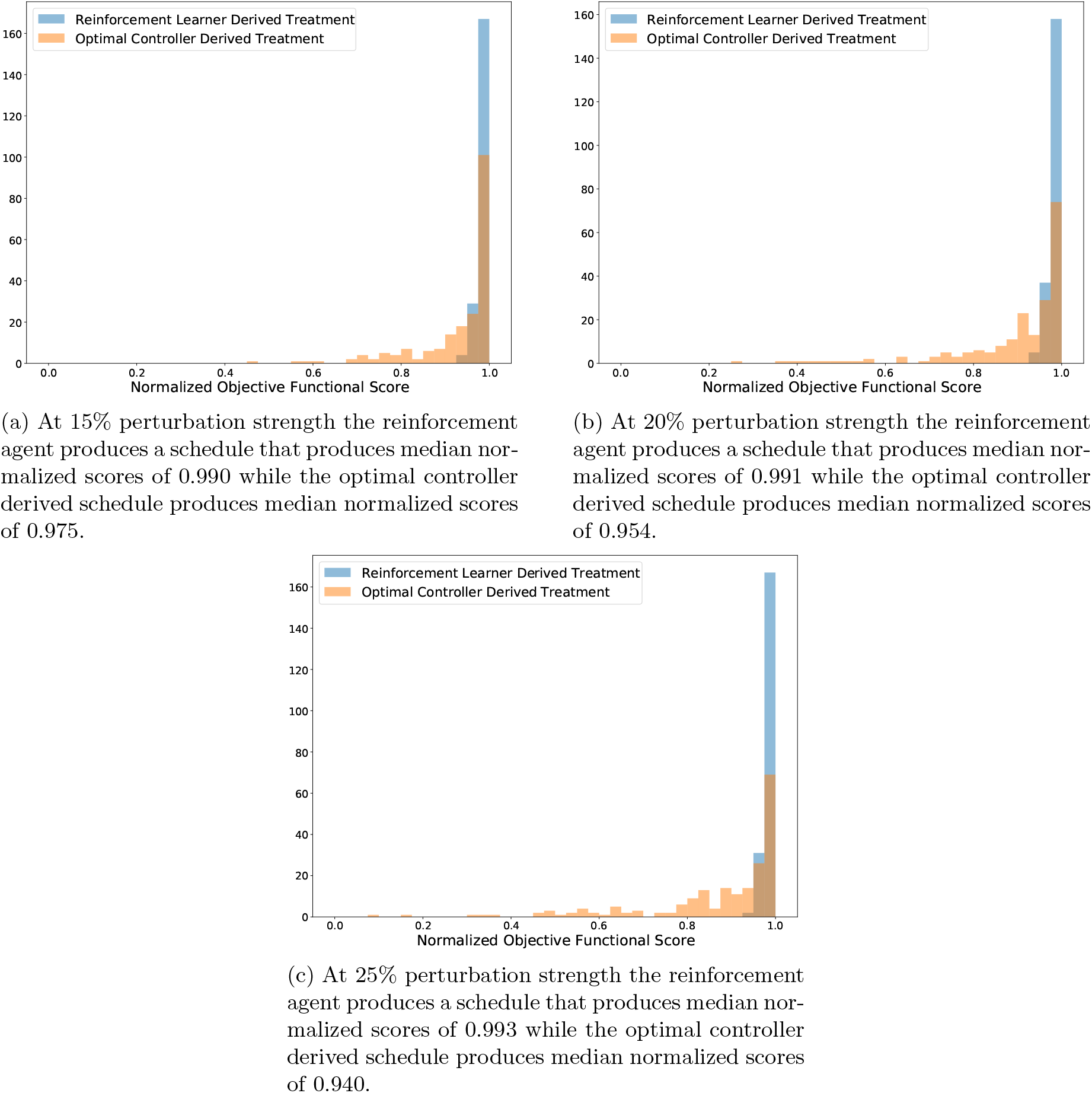
Histograms of the scores achieved by the various agents on the 200 testing virtual patients. Bin sizes were chosen to correspond to 0.025. In particular, the reinforcement learning agent is much more robust toward perturbation in parameter values, consistently producing dosing schedules scoring within 7.5% of the theoretical maximal score.

### 4.2 Contrasting a nominal reinforcement learning agent with a nearest neighbour interpolated optimal controller

The results of Section 4.1 indicate that the reinforcement learning agent produces schedules that are more robust to perturbations in the unknown parameters. We noted that the reinforcement learning agent is capable of customizing these schedules for each individual patient, not by measuring the patient specific parameters directly, but by customizing the response via a more easily measurable metric. In this section, we consider a different training process that allows the optimal controller agent a comparable level of customization.

For this comparison, we kept the reinforcement learning agent exactly the same as in Section 4.1: the agent was trained on only the nominal parameter set and was kept blind to the testing and training virtual patients. For the optimal controller comparison, we begin by solving the discrete optimal control problem on all 50 training virtual patients from Section 3.2. We then log the state vector from Eq. (7) for each timestep of treatment. For each of the 200 testing patients at each time step *t*_*i*_ we calculated the state vector 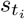, we then applied a chemotherapeutic dose by consulting the table of states from the training patients at time *t*_*i*_. The dose was selected from the training patient whose state vector was closest to the current testing patient state vector (where distance was measured by the Euclidean metric). The result of this nearest training neighbour optimal controller (NTNOC) was an agent that could also customize chemotherapeutic dosing strategies for each of the 200 testing virtual patients based off of knowledge gained by traversing the training virtual patient space. In contrast, the reinforcement learning agent it is being compared to only ever interacted with a differential equation environment parameterized by the nominal parameter set. Ostensibly, more distribution level information is directly afforded to the NTNOC than was afforded to the reinforcement learning agent. The reinforcement learning agent is only able to customize treatment strategies based off the states learned by providing non-optimal doses to the nominal virtual patient during training. The results of this comparison are presented in Figure 9. In particular, we note that the same general trend from Figure 7 is repeated: namely, as the perturbation strength increases the relative performance of the reinforcement learning agent also increases. However, in contrast to Figure 7, we note that at the 15% level the nearest training neighbour optimal controller outperforms the reinforcement learning agent (with a one-sided Wilcoxon signed-rank test p-value on the order of machine precision). Indeed, the mean value of the differences plotted in Figure 9a occurs at approximately −0.008, indicating that, in a mean value sense, the nearest training neighbour optimal controller produces strategies that are 0.008 points closer to the optimal score of 1 than the scores of the schedules produced by the reinforcement learning agent.

**Figure 9:**
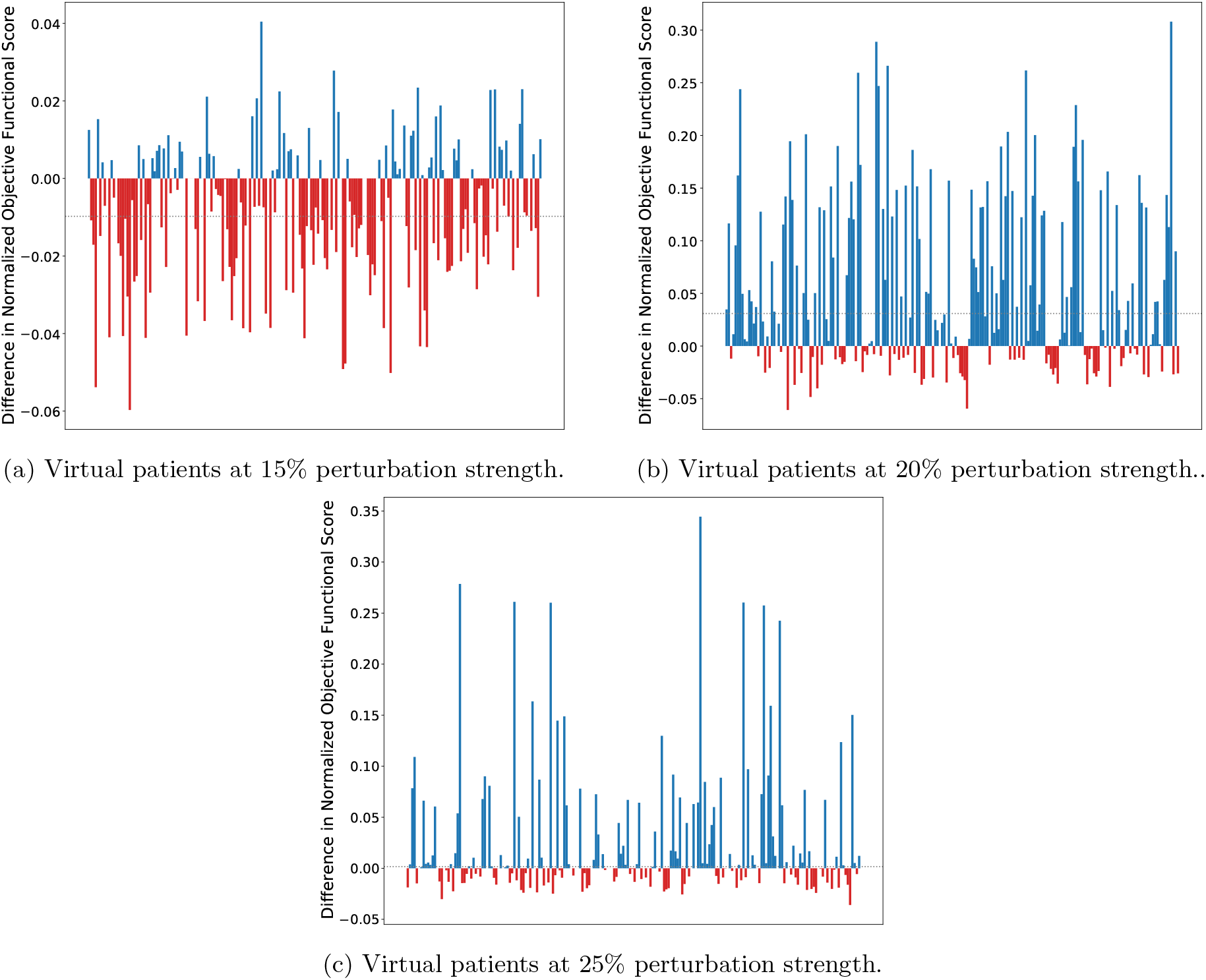
Bar plots of the difference between the scores obtained by the reinforcement learner derived policy and the scores obtained by the nearest training neighbour optimal controller on all test virtual patients. Testing patients where the reinforcement learner outperformed the optimal controller are marked in blue and patients where the optimal controller outperformed the reinforcement learner are marked in red. The dotted grey lines in each plot indicate difference in the median values of the scores obtained by the reinforcement learning agent and those obtained by the nearest training neighbour optimal controller.

Again we compare the distributions of scores with the one-sided Wilcoxon signed-rank test, though for the case represented in Figure 9a, we consider the alternative hypothesis to be that the nearest training neighbour optimal controller produces schedules with higher median normalized score than that of the reinforcement learning agent. We found at the 15% perturbation strength level a Wilcoxon statistic of 4997 corresponding to a *p*-value on the order of 10^−9^. Hence we reject the null hypothesis and conclude that, at the 15% perturbation level, that the NTNOC produces chemotherapeutic schedules with higher median normalized score than those produced by the reinforcement learning agent. For the cases represented in Figures 9b and 9c we consider a different alternative hypothesis: namely that the reinforcement learning agent produces chemotherapeutic schedules with higher median normalized score than those produced by the NTNOC. Then, at the 20% perturbation strength level we found a Wilcoxon statistic of 15706 corresponding to a *p*-value on the order of 10^−13^, and at the 25% perturbation strength level we found a Wilcoxon statistic of 11051 corresponding to a *p*-value on the order of 10^−3^. In these two cases we reject the null hypothesis and conclude the reinforcement learning agent produces chemotherapeutic schedules with higher median normalized score on perturbed patients than the NTNOC. In this situation, the nearest training neighbour optimal controller is able to produce schedules more competitive with the reinforcement learning agent than those produced by the nominal optimal controller. In the 15% case, the NTNOC outperforms the reinforcement learning agent by a small margin (difference in median scores of 0.009 in favour of the NTNOC) whereas the reinforcement learner outperforms the NTNOC in the 20% case (difference in median scores of 0.031 in favour of the reinforcement learner) and the 25% case (difference in median scores of 0.002 in favour of the reinforcement learner). While the NTNOC produces more robust schedules for small perturbations, such schedules seem to only slightly outperform the schedules produced by the reinforcement learning agent. In contrast, for medium perturbations around 20% and 25%, the reinforcement learning agent outperforms the NTNOC.

In Figure 10 we concern ourselves with, once again, examining the histograms of the scores of these two agents. As before, we note that the reinforcement learner derived schedules are robust to these perturbations in patient specific parameter values, which is the source of the success in Figure 9b and Figure 9c. However, in contrast to Figure 8, we note that the nearest training neighbour optimal controller produces schedules whose scores produce a histogram that is less diffuse than that produced by the nominal optimal controller (standard deviations of (0.009, 0.07, 0.06) for the nearest training neighbour optimal controller at the 15%, 20%, and 25% perturbation strength compared to standard deviations of (0.09, 0.14, 0.17) for the nominal optimal controller and (0.01, 0.01, 0.01) for the reinforcement learning agent). The end result is, by allowing the optimal controller access to more distribution-level data, it is capable of customizing schedules in a way that is more robust to perturbations in the patient specific parameters. However, for sufficiently high perturbations in the strength of the parameters, these personalized schedules are still less optimal than the personalized schedules produced by the reinforcement learning agent. Again, we note that the success of the reinforcement learning agent is due to the increased diffusivity of the distribution of scores obtained by the NTNOC as perturbation strength increases as contrasted with the more stable distribution of reinforcement learning agent derived scores under the same perturbation strengths.

**Figure 10:**
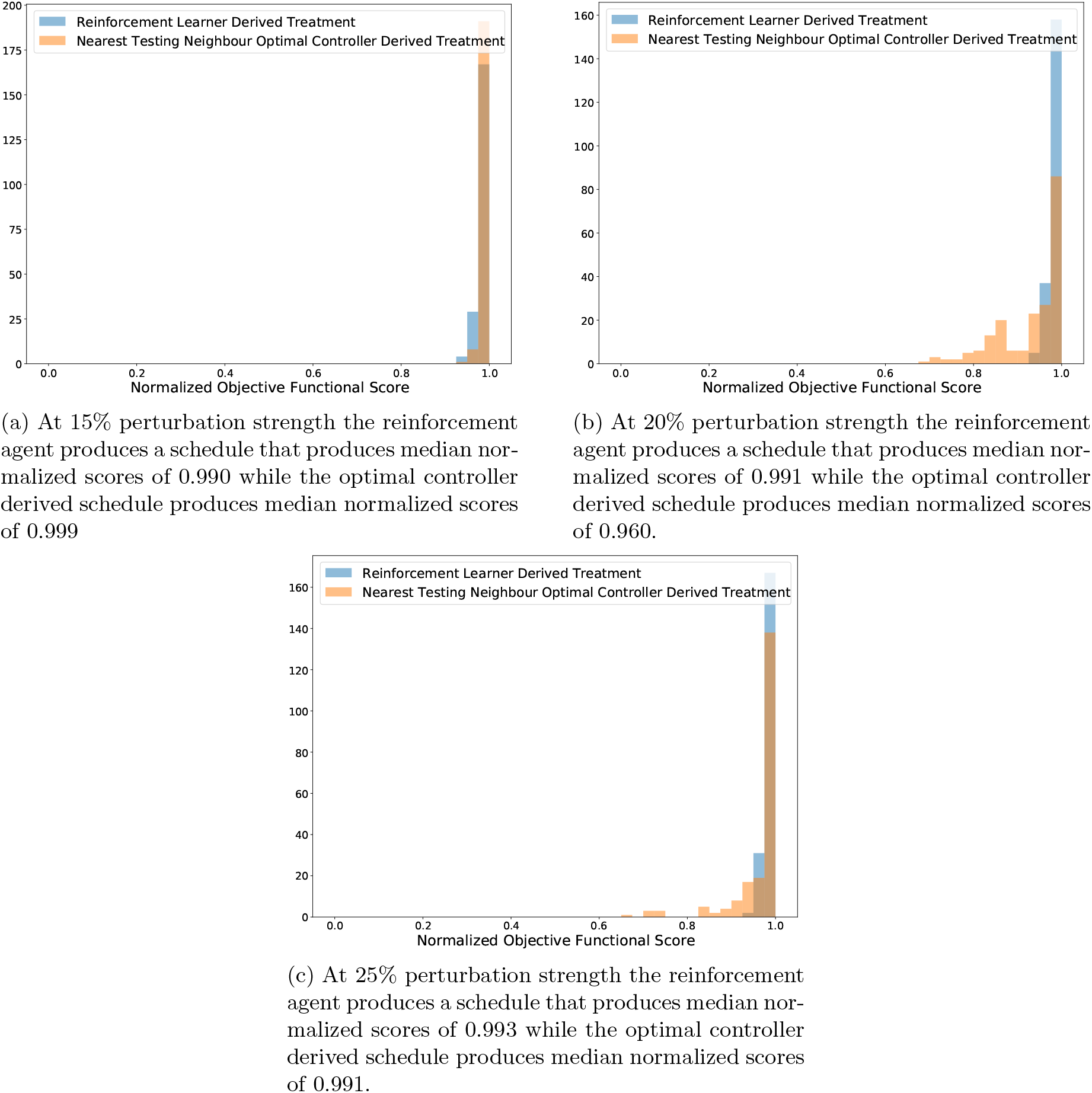
Histograms of the scores achieved by the various agents on the 200 testing virtual patients. Bin sizes were chosen to correspond to 0.025. In particular, while the reinforcement learning agent is more robust towards perturbation in parameter values at the 20% and 25% perturbation strength, the nearest neighbour optimal controller produces schedules within 5% of the theoretical maximum at the 15% perturbation level.

## 5 Conclusion

In summary, we examined a model of breast cancer, ovarian cancer, and bone marrow density under treatment by a chemotherapeutic for which the continuous time optimal control has been analytically derived. We discretized the optimal control problem of chemotherapeutic dosing schedule under the objective functional in Eq. (5) to apply different doses every day with dose strength discretized to be in 0 to 1 inclusive by steps of size 0.1. By solving this discretized problem on 200 testing virtual patients, we were able to establish ground truth levels for theoretical maximal objective functional scores. We then contrasted a reinforcement learning agent trained on the nominal parameter set with a traditional optimal controller on the nominal parameter set. We noted that since the reinforcement learning agent trains a fixed policy, it can customize the corresponding dose schedule to testing virtual patients, even when the patient-specific parameterization of the differential equation environment from Eq. (1) is unknown, by leveraging more data that is easier in practice to collect. In this case, this meant providing the reinforcement learning agent with a window of relative bone marrow density mass (relative to before the treatment process begun). We noted that the reinforcement learning agent produces schedules that are closer to the theoretical optimum in a mean sense when measured against unknown patients who differ from the nominal parameter set by 15%, 20%, and 25%. In particular, we note that as the strength of perturbation increases, the net benefit of using the reinforcement learning derived schedules also increases. Moreover, as the perturbation strength increases, the collection of normalized optimality scores stay clustered between 0.925 and 1. In contrast, as the perturbation strength increases, the collection of normalized optimality scores for the optimal controller become more diffuse.

We also allowed the optimal controller derived schedules to leverage the longitudinal relative bone marrow density information by training 50 such optimal controllers on perturbed parameter values that were treated as known. When we compared this nearest training neighbour agent to the reinforcement learning agent, we discovered that the reinforcement learning agent still outperformed the other agent at the 20% and 25% perturbation strength level, but at the 15% perturbation strength level the nearest training neighbour agent was more optimal. However, this nearest training neighbour optimal controller was still prone to reduced performance level and a more diffuse histogram of dose-schedule scores at the higher perturbation levels, something that we did not observe in the reinforcement learning agent.

We conclude by noting that reinforcement learning provides an agent that can be used to personalize dosage schedules in the absence of patient specific parameter data in a manner that is not prone to wild fluctuations (as evidenced by the tight histograms of Figure 8 and Figure 10) and did so by only requiring the mean values of the patient specific parameter distributions. In contrast, an optimal control derived agent could be improved to allow personalization of dosing schedule as well, but at the cost of requiring more samples from these patient specific distributions and the end result was still prone to fluctuations in schedule optimality.

While this particular study was focused on a situation where little patient data was used (outside of the data used to determine the nominal parameters from Table 1) one could also extend this work by allowing a reinforcement learner to learn directly from patient data, since the environment a reinforcement learning agent interacts with is effectively a black box. This method described is general and can be used for optimizing schedules for other treatments as well (i.e. radiotherapy fractions or immunotherapy treatment schedules). Moreover, this study was focused on a particular model in a mathematical oncology context, however we believe this result can be applied to other mathematical models used in cancer research and can also be extended easily to other mathematical biology contexts, or into any context wherein one needs to create a control for a system where high level distribution information about the parameters is known, but particular parameter values are unknown or prohibitively difficult to ascertain.

## Acknowledgment

This work was supported by the Canadian Institutes of Health Research (CIHR).

## Supplementary Material

In Figure 11 we present a visualization of the virtual patients used for testing of all three algorithms and the training virtual patients used for training of the NTNOC algorithm. These figures demonstrate the parameter values chosen via Latin hypercube sampling, as described in Section 3.2, for the four non-zero bone marrow parameters from Table 1 required to parameterize Eqn. 5.

In Figure 12 we present the trajectories of the bone-marrow and the control obtained via the objective functional in Eqn. 6.

**Figure 11:**
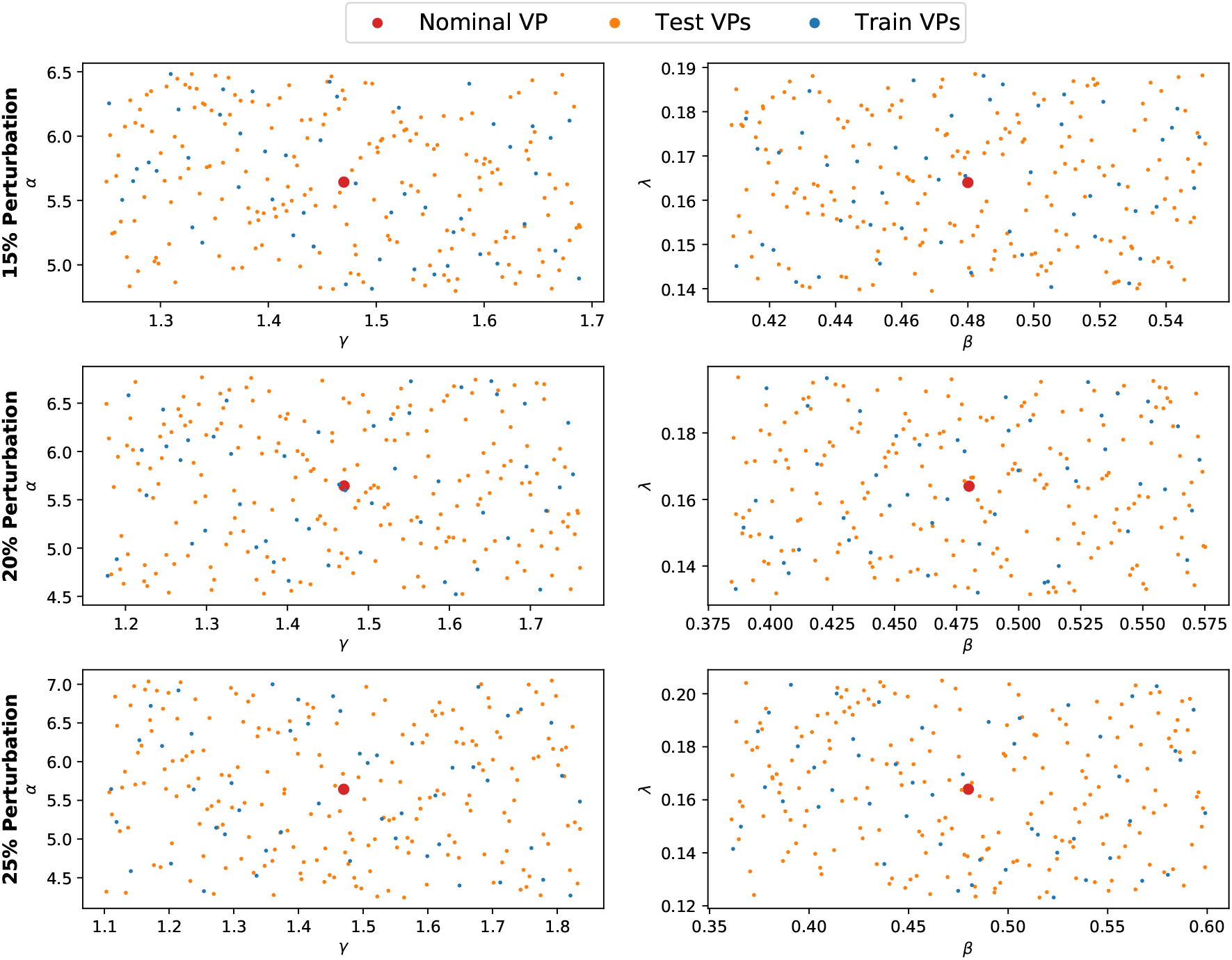
Visualization of the 4 non-zero virtual patient parameters for various perturbation strengths. The orange dots indicate the 200 virtual patients used in testing, the blue dots the 50 virtual patients used in training, and the red dot indicates the location of the nominal virtual patient.

**Figure 12:**
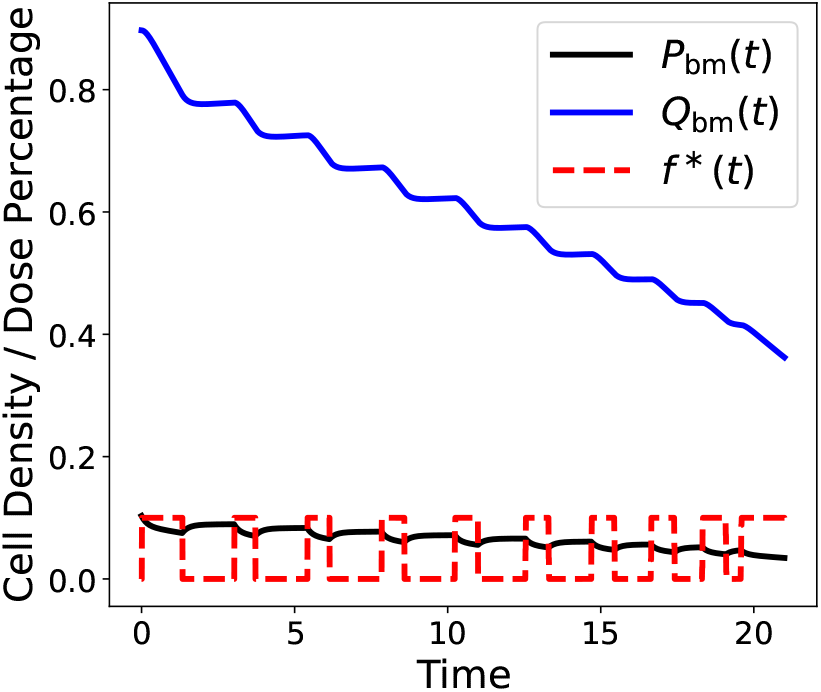
A plot of the Eqn. 1 parameterized for bone-marrow under the optimal control achieved by maximizing the objective functional in Eqn. 6 with *b* = 2. Solved numerically using IPOPT in GEKKO [5, 18].

Figure 13 demonstrates various dosing schedules on testing patients. We plot the schedules obtained via employing the reinforcement learning agent trained on the nominal parameter set, the mean optimal controller derived treatment, the NTNOC treatment, and the optimal treatment (which was treated as unknown) acquired by solving the optimal control problem on each particular testing patient (achieved by treating the model parameters as known and employing the APOPT algorithm as implemented in GEKKO [5, 15]).

**Figure 13:**
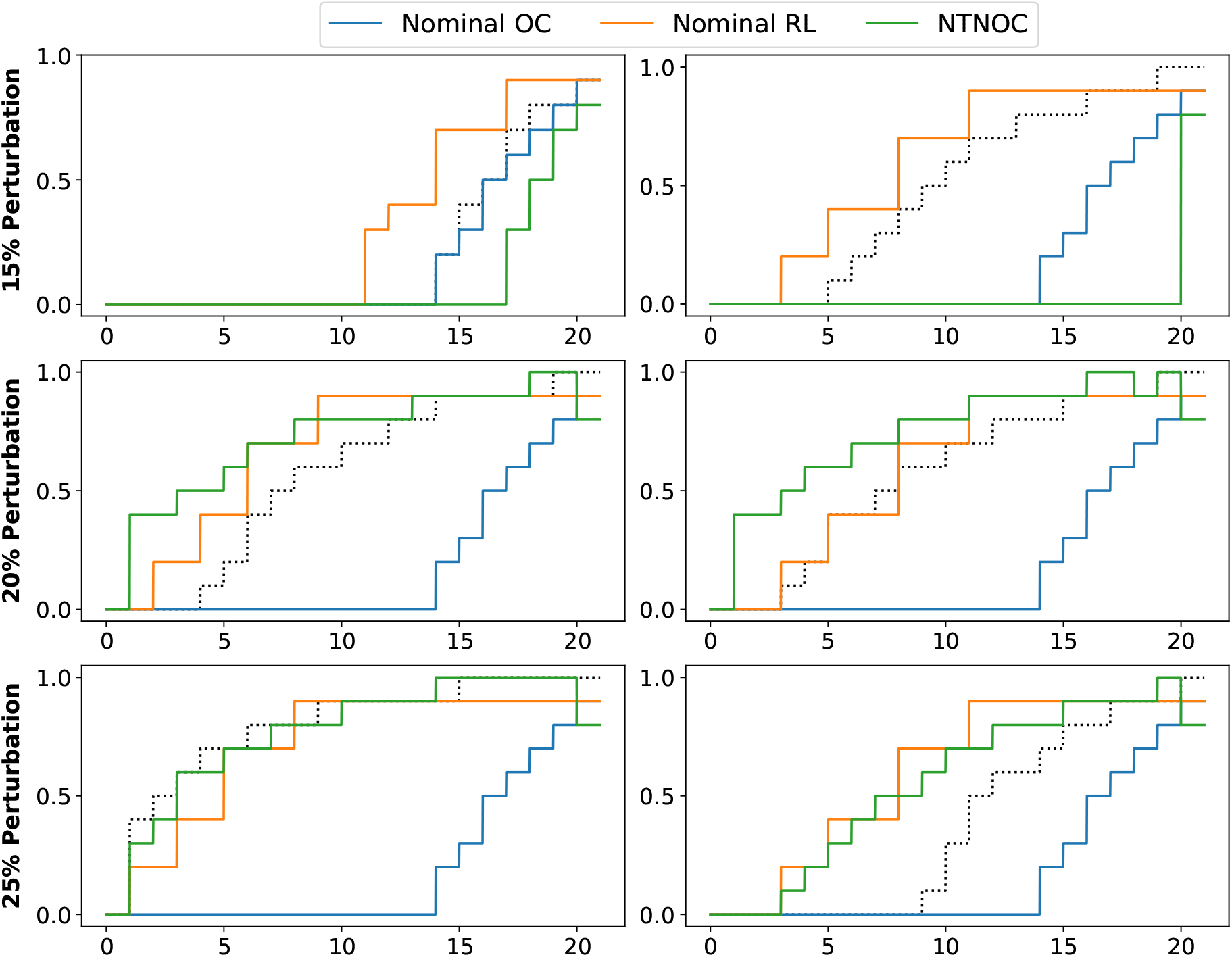
Various chemotherapy dosing schedules obtained on various virtual patients by the three blind methods. In dotted black lines, we also present the theoretically optimal schedule for each patient as a comparison.

